# The spread of antibiotic resistance is driven by plasmids amongst the fastest evolving and of broadest host range

**DOI:** 10.1101/2024.07.23.604842

**Authors:** Charles Coluzzi, Eduardo PC Rocha

**Affiliations:** Institut Pasteur, Université Paris Cité, Microbial Evolutionary Genomics, CNRS UMR3525, 75724 Paris, France

## Abstract

Microorganisms endure novel challenges for which other microorganisms in other biomes may have already evolved solutions. This is the case of nosocomial bacteria under antibiotic therapy because antibiotics are of ancient natural origin and resistances to them have previously emerged in environmental bacteria. In such cases, the rate of adaptation crucially depends on the acquisition of genes by horizontal transfer of plasmids from distantly related bacteria in different biomes. We hypothesized that such processes should be driven by plasmids amongst the most mobile and evolvable. We confirmed these predictions by showing that plasmid families encoding antibiotic resistance are very mobile, have broad host ranges, while showing higher rates of homologous recombination and faster turnover of gene repertoires than the other plasmids. These characteristics remain outstanding when we remove resistance plasmids from our dataset, suggesting that antibiotic resistance genes are preferentially acquired and carried by plasmid families that are intrinsically very mobile and plastic. Evolvability and mobility facilitate the transfer of antibiotic resistance, and presumably of other phenotypes, across distant taxonomic groups and biomes. Hence, plasmid families, and possibly those of other mobile genetic elements, have differentiated and predictable roles in the spread of novel traits.

## Introduction

Bacteria adapt quickly to environmental challenges, including stress, antibiotic treatment, or prophage predation. This is facilitated by their ability to acquire genes by horizontal transfer using mobile genetic elements (MGEs) [1, 2]. These elements can be mobilized by conjugation or phage transduction and remain in cells as plasmids (extrachromosomal elements) or integrated in the chromosome [3]. Many potentially adaptive accessory traits are carried by conjugative elements or other MGEs mobilized by them or by phages and their satellites [4]. One might expect that such adaptive genes would tend to get fixed in the novel genome. Yet, functions under balancing selection, whose fitness value varies with time, space, or frequency in the population [5], may be maintained in MGEs because they are constantly gained and lost. Functions may also remain in plasmids because gene expression and evolutionary rates may be modulated by plasmid copy number [6, 7]. In all these cases, there is either an intermittent or a constant pressure to maintain a trait in a plasmid that circulates in a community. But certain functions may be predominantly found in MGEs because selection for the trait is recent and genes encoding such a trait were not initially available in the community. In such cases, genes are predominantly in MGEs because they are just arriving in the local community and MGEs are their vehicles.

An example of the importance of MGE-mediated horizontal gene transfer results from the challenge imposed by the introduction of antibiotic therapy to control human-related bacteria in the last few decades [8]. Under such an unprecedented challenge, adaptation required either functional innovation, which is often a slow process, or the acquisition of antibiotic resistance genes from bacteria inhabiting natural environments where antibiotics have been present for a long time [9]. Indeed, horizontal gene transfer, especially of plasmids, is driving the rapid spread of antimicrobial resistance (AMR) genes in the last decades [10]. Importantly, AMR is driven by plasmids that lacked these genes 70 years ago [11]. Conjugative plasmids were shown to disseminate resistance to penicillins, aminoglycosides, sulfonamides and last-resort antibiotics such as colistin or carbapenems [12–15]. They are also frequently associated with the spread of pandemic multidrug-resistant high-risk clones such as *Escherichia coli* sequence type 131 and *Klebsiella pneumoniae* sequence type 258 [16, 17]. Integrative conjugative elements and phage-plasmids also contribute to the spread of AMR genes [18, 19]. This raises the question of which MGE characteristics favor the spread of novel traits.

We hypothesize that two characteristics are crucial for gene transfer between distantly related bacteria living in different biomes: the ability of MGEs to acquire, exchange and deliver genes (genetic plasticity, here considered as evolvability by large genetic changes) and their ability to transfer to distant hosts (mobility and especially host range). The first may result from allelic exchanges between homologous genes in MGEs (homologous recombination). Alternatively, genetic exchanges may be mediated by recombinases, including transposable elements and integrons [20, 21]. A recent study reported that 77% of the plasmids carrying

AMR genes have insertion sequences [22], and many studies described intragenomic transfer of resistance genes in composite transposons and integrons [23, 24]. For instance, the colistin resistance gene *mcr-1* is present in many different plasmid types largely because of its association with the ISApl1 transposon [14]. Recombinases facilitate the translocation of AMR genes in the environmental bacteria encoding antibiotic resistance, their subsequent transfer between plasmids, and their potential future integration in chromosomes [25]. Of note, plasmids have by far the highest densities of transposable elements among bacterial replicons. Mobility is also essential for genetic transfer and host range is thus expected to contribute to explain the preferential presence of AMR genes in some plasmids. Broad host range plasmids are more likely to transfer from distantly related environmental bacteria. These variables may combine to produce particularly effective vectors of AMR. For example, the broad host-range plasmid pNDM-1_Dok01 partly responsible for the spread of β-lactamase resistance in *E. coli* ST38 carries the New Delhi metallo-β-lactamase (NDM-1) in a composite transposon very similar to the ones of distantly related environmental bacteria [26].

Comparisons between antibiotic resistance plasmids often reveal a quick pace of change for these elements [27], whereas other plasmids are remarkably stable [28]. This led us to hypothesize that plasmids carrying antibiotic resistance in current nosocomial bacteria are among those that are most plastic (in terms of rates of allelic exchanges and pace of change of gene repertoires), mobile (able to transfer or be transferred horizontally), and with broader host ranges. These characteristics would contribute to explain why transfer of a novel resistance occurs rapidly after the introduction of the antibiotic on the market [29]. Our hypothesis might have broader implications: when an unprecedented challenge leads to strong selection for novel traits that are absent in a community, their acquisition from other communities will be driven by MGEs particularly plastic and with broad host ranges. To test our hypothesis, we focus specifically on antibiotic resistance because of its clinical relevance and data availability. We focus on plasmids because they are the key spreaders of AMR and because they are easy to identify and delimit in completely assembled genomes. More precisely, we identified the plasmids carrying and lacking antibiotic resistance genes, clustered them in plasmid taxonomic units (PTUs), and analyzed them in terms of mobility, host range, homologous recombination, and gene turnover. Finally, we integrated these results to rank their relevance and to understand if these plasmids are more plastic because they carry resistance genes or because they are part of particularly plastic plasmid families.

## Materials and Methods

### Plasmid dataset

The dataset used in this study consists of 32,798 complete genomes from 8,345 species retrieved from NCBI RefSeq database of high-quality complete non-redundant prokaryotic genomes (https://ftp.ncbi.nlm.nih.gov/genomes/refseq/, last accessed in May 2023, [30]). These genomes contain 38,057 plasmids. To avoid the misidentification of secondary chromosomes as plasmids, 996 plasmids larger than 500 kb were excluded (Harrison et al. 2010; Smillie et al. 2010). The complete list of 37,061 plasmids can be found in supplementary table S1.

### Coding sequence prediction

We have previously observed that some automatic annotations mis-annotate some genes in MGEs, notably the relaxase, which is essential to study plasmid mobility. To ensure that sequence annotation is homogeneous and appropriate for our purpose, all plasmid sequences were re-annotated using prodigal *2.6.3* [31] with the *‘-p meta’* option to obtain the putative protein coding genes.

### Plasmid mobility prediction

Conjugative systems were detected using MacSyFinder *v2.07rc* [32] with the ‘Plasmids’ models of the CONJScan module. Briefly, MacSyFinder uses HMM protein profiles and a set of rules (which are defined in the models) about their presence and proximity. MacSyfinder was used with default parameters and CONJScan models were ran all at once with the parametters ‘-all -ordered -circular’. The plasmids were then classed in relation to the conjugation genes they contain (as in [33]). The plasmids encoding a complete mating pair formation system and a relaxase were classed as conjugative (**pCONJ**). The remaining plasmids that encode a relaxase were classed as mobilizable plasmids (**pMOB**). Plasmids that are none of the above but have an oriT and may thus be mobilized by a relaxase and a MPF encoded in *trans* were classed as **pOriT**. These origins of transfer were identified using *blastn* (*v2.09.0+*) [34] with parameters *-task blastn-short -evalue 0.01* filtered at identity >80% and coverage >80% and a dataset of 91 sequences of different origins of transfer [35].

Plasmids were classed as phage-plasmids (**P-P**) as in [36]. The remaining plasmids were considered as putatively non-transmissible by conjugation (**pNT**).

### Detection of antibiotic resistances, integrons and insertion sequences

Plasmid protein sequences were screened for the genes conferring antibiotic resistance to the 31 AMR drug classes detected by AMRFinderPlus *v3.11.14* with default parameters (identity > 90%, e-value < 1e-20, coverage > 50%) [37] and the database version *2023-04-17.1*. Plasmids encoding at least one antibiotic resistance gene were considered AMR+ plasmids. Through the text, AMR classes and AMR genes correspond to “AMR classes” and “AMR gene symbol” as defined by AMRFinderPlus.

Insertion sequences were detected using ISEScan *v1.7.2.3* [38] with default parameters and the ‘--removeShortIS’ option.

Integrons were detected using IntegronFinder *v2* [39] with the ‘–local-max’ and ‘—circ’ options.

### PTU and host-range predictions

The Plasmid Taxonomic units (PTU) and host-Range were assigned to each plasmid with COPLA *v1.0* and default parameters [40]. The host-Range indicates the ability of a PTU to colonize different bacterial species. Each PTU has a host-range grade that reflects the diversity of bacterial species in which the plasmids within that PTU have been found. Host-range grades vary between I and VI, the higher the grade, the broader the host range, where grades correspond to PTUs:

- Grade I: Plasmids of the PTU are found in strains of the same species.
- Grade II: Plasmids of the PTU are found in species of the same genus.
- Grade III: Plasmids of the PTU are found in genera that belong to the same family.
- Grade IV: Plasmids of the PTU are found in different taxonomic families of the same order.
- Grade V: Plasmids of the PTU are found in different orders of the same class.
- Grade VI: Plasmids of the PTU are found in different classes of the same phylum or even beyond, in different phyla.

Grades V and VI were merged to have enough counts for statistical tests.

### PTU pangenome and core-genome construction and phylogeny inference

The pangenome and core-genome of each PTU were computed using PanACoTA *v.1.3.1* [41]. Briefly, the pangenome was constructed by clustering all protein sequences in the set of genomes using MMseqs2 (protein identity >80% [42]) We retrieved the core-genome from each pangenome. It consists of the gene families present in exactly one copy in at least 80% of all plasmids.

The multiple sequence alignments of the families of the core genes were computed using the ‘align’ PanACoTA *v1.3.1* module. Briefly, the protein sequences of the core genes were aligned with MAFFT *v7.467* (-auto parameters) [43] then back-translated to nucleotide alignments (*i.e.*, each amino acid was replaced by the original codon) and concatenated. The phylogenetic inference was done from the resulting multiple alignments using IQ-TREE *2.0.6* [44] with the ultra-fast bootstrap option (-bb 1000 bootstraps) and with the best fitting model estimated using *ModelFinder Plus* (-MFP) [45]. The best model for each tree was determined according to the BIC criterion. To obtain robust phylogenetic reconstructions, we followed [46] and removed from the phylogenetic analyses the plasmids encoding less than 50% of the core genes of a PTU.

The average tip-to-root distance for each phylogenetic tree was computed using the “get_distance()” function of ETEToolkit *v3.1.2* [47]

### Rarefaction curve and Heaps’ law fit

Rarefaction curves for the pangenome of each PTU were computed using the package vegan for R, v2.5.6 (https://CRAN.R-project.org/package=vegan) on the presence/absence matrixes of gene families obtained by PanACota. As rarefaction curves require a large sample size to be accurate [48], we computed them only for PTUs with more than 30 plasmids. The Heaps’ law was fitted to each rarefaction curves [49].

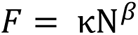

Where F is the number of pangenome families and K and β are empirical parameters that were extracted using the ‘optimize.curve_fit’ method from scipy *v.11.4* for python [50].

### Assessment of congruence between phylogenetic trees

The Robinson-Foulds symmetric distance between two trees corresponds to the number of splits or fusions that are required to transform the topology of one tree into another. The normalized Robinson-Foulds symmetric difference metric (nRF) corresponds to the Robinson-Foulds symmetric distance [51] divided by the value of the theoretical maximum of the Robinson-Foulds distance. The nRF varies between 0 and 1, where pairs of trees with nRF=0 have identical topologies and those with nRF=1 have no common splits. For each PTU with a phylogenetic tree based on more than 1 core gene multiple alignment, we computed the mean nRF between the tree constructed from all concatenated core genes of the PTU and the tree constructed from each individual gene within the core genome (**concatenate-based comparisons**). This allows to measure the degree of incongruence between each individual core gene and the set of all core genes. High values of nRF are thus an indication of recombination. Of note, such tests are usually made with trees that do not include the genes being tested, e.g. robust phylogenies produced by external data. Yet, such trees are unavailable for plasmids. Given the recombination rates we observe, they are probably impossible to identify in many PTUs thus also precluding the use of recombination detection methods based on the bacterial clonal frame [52]. The use of the focal gene in the comparison (as part of the genes that were used to build the core genes phylogeny) renders the method conservative, i.e. it will tend to produce lower rates of incongruence. To assess this potential problem, we also computed the nRF between the tree of every gene of the core genome of the PTU (**all-pairs comparisons**, self-comparisons are omitted, i.e. tree of the gene A is not compared to itself). In this case there is no assumption of a core gene single phylogeny. These scores were computed using *ete-compare* method from the ETEToolkit *v3.1.2* [47]. To ensure only the robust parts of the trees were considered, all branches with a bootstrap support inferior to 75 were discarded, with the ‘min_support_src’ and ‘min_support_ref’ options.

### wGRR calculation

We estimated the weighted gene repertoire relatedness (wGRR) values between each pair of plasmids in our dataset using the wGRR software (https://gitlab.pasteur.fr/jgugliel/wgrr) with default parameters as described in [53]. Concretely, we identified plasmid proteins that are significantly similar (identity ≥ 35%, coverage ≥ 50%) among all pairs of plasmid sequences with MMseqs2 *search v. 9-d36de* with the parameter “sensitivity 7.5 (-s 7.5) and “max-seqs 100000” (to retrieve all possible hits). We then identified the best bidirectional hits (BBH) between pairs of plasmids which correspond to couples of proteins that are reciprocal best matches. Finally, we use the BBH to calculate the wGRR.

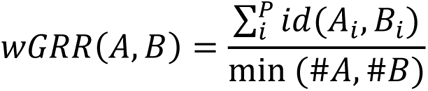

Where A*_i_* and B*_i_* are the *ith* BBH pair of P total pairs, id(A*_i_*, B*_i_*) is the identity between the pair, and min(#A, #B) is the number of genes of the plasmid having the fewest. The wGRR varies between 0 and 1. A wGRR close to 1 means that both plasmids are very similar (all genes in a plasmid have very similar best bi-directional hits in the other plasmid) and a wGRR of 0 means that plasmids have no homologs.

For each PTU, we computed the median and mean wGRR values between every possible pair of plasmids within the PTU. A high mean wGRR value means that most of the plasmids in the PTU are closely related and a small mean wGRR value means that most plasmids in the PTU are more distantly related.

### Statistics

ANOVA, χ^2^ tests, ordinal and nominal logistic regressions were performed using the “Fit model” module of JMP *v17.* All others statistical tests were performed within R (*v. 4.2.3*)

## Results

### Plasmid groups are heterogeneously enriched in ARGs

We retrieved all plasmids sequenced within complete genomes of the non-redundant database RefSeq of the NCBI to study their evolution. We excluded the few very large elements that could be secondary chromosomes or chromids (> 500 kb) (Figure S1, Table S1), re-annotated the remaining 37,061 plasmids, and screened them for antibiotic resistance genes. We used conservative methods to focus on genes with high similarity to known antibiotic resistance genes in nosocomial pathogens (identified with AMRFinderPlus). About a quarter of them (9,384) encode at least one antimicrobial resistance gene (Figure 1 A/B). These plasmids are 60% larger than the others (Figure 1C). We analyzed in detail the 392 unique antibiotic resistance genes (determined as ‘gene symbol’ by AMRFinderPlus) present in total 46,472 times in these plasmids (Table S1). These genes conferred resistance to 31 drug classes (determined as ‘AMR class’ by AMRFinderPlus). The most common antibiotic resistance classes found in the dataset target beta-lactams (6,032 plasmids), aminoglycosides (5,250 plasmids) and sulfonamides (3,725 plasmids) (Figure 1D). These genes often co-occur with those of other AMR classes (Figure 1D). We found 895 different combinations of AMR classes in plasmids, some of which are more often found in combination than alone (Figure 1D). Accordingly, plasmids with antibiotic resistance genes have a median of three different AMR classes.

**Figure 1:**
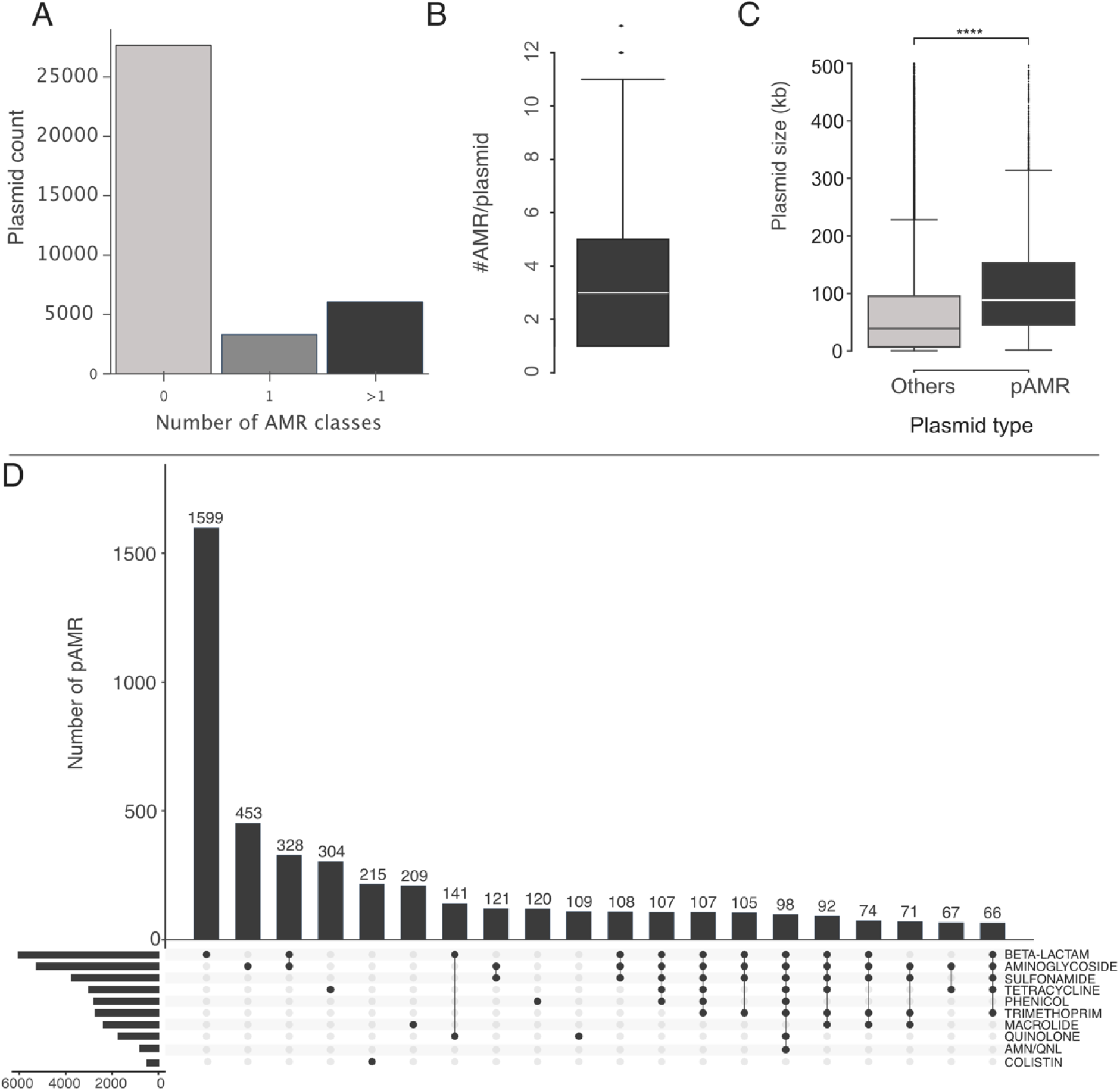
Antibiotic resistance (AMR) classes in plasmids. **A**. Number of plasmids with one or more AMR classes. **B**. Number of AMR classes per plasmid carrying at least one resistance gene. **C**. Distribution of the size of plasmids encoding AMR ("pAMR") or not ("Others"). The horizontal bar in boxplots indicates the median value, and lower and upper hinges correspond to the first and third quartiles. The whiskers extend from the hinge to 1.5 times the range between the first and third quartile. Data beyond these values are shown as dots. Statistical significance (Mann-Whitney U): **** P ≤ 0.0001. **D**. Upset plot representing the co-occurrence of the 10 most abundant AMR classes and the 20 most common AMR classes combinations in the dataset. The horizontal bar plot on the bottom left shows the frequency of each AMR class.

To assess if different combinations of antibiotic resistance classes could often be found in closely related plasmids, we identified the plasmid taxonomic units (PTU) of the 37,061 plasmids. PTUs are cohesive groups of plasmids that share high average nucleotide identity and approximately similar size. Almost half (17,314) of the plasmids and two thirds (6,321) of those containing resistance genes could be assigned to one of the previously described 355 PTUs (Table S1, [40]). 223 (63%) PTUs lacked plasmids with antibiotic resistance genes. Among the others, the plasmids encoding resistance genes had widely variable frequencies, some of them only containing such plasmids (Figure 2, Figure S2, Table S2). Most of these PTUs contained several combinations of AMR classes. The PTU-HI2 (including pSTM6 and relatives) is an extreme case: 98% of its plasmids have resistance genes and these plasmids harbor 197 different combinations of AMR classes (Figure S3). The high diversity observed among closely related plasmids (within the same PTU) suggests that resistance genes were acquired in multiple events.

**Figure 2:**
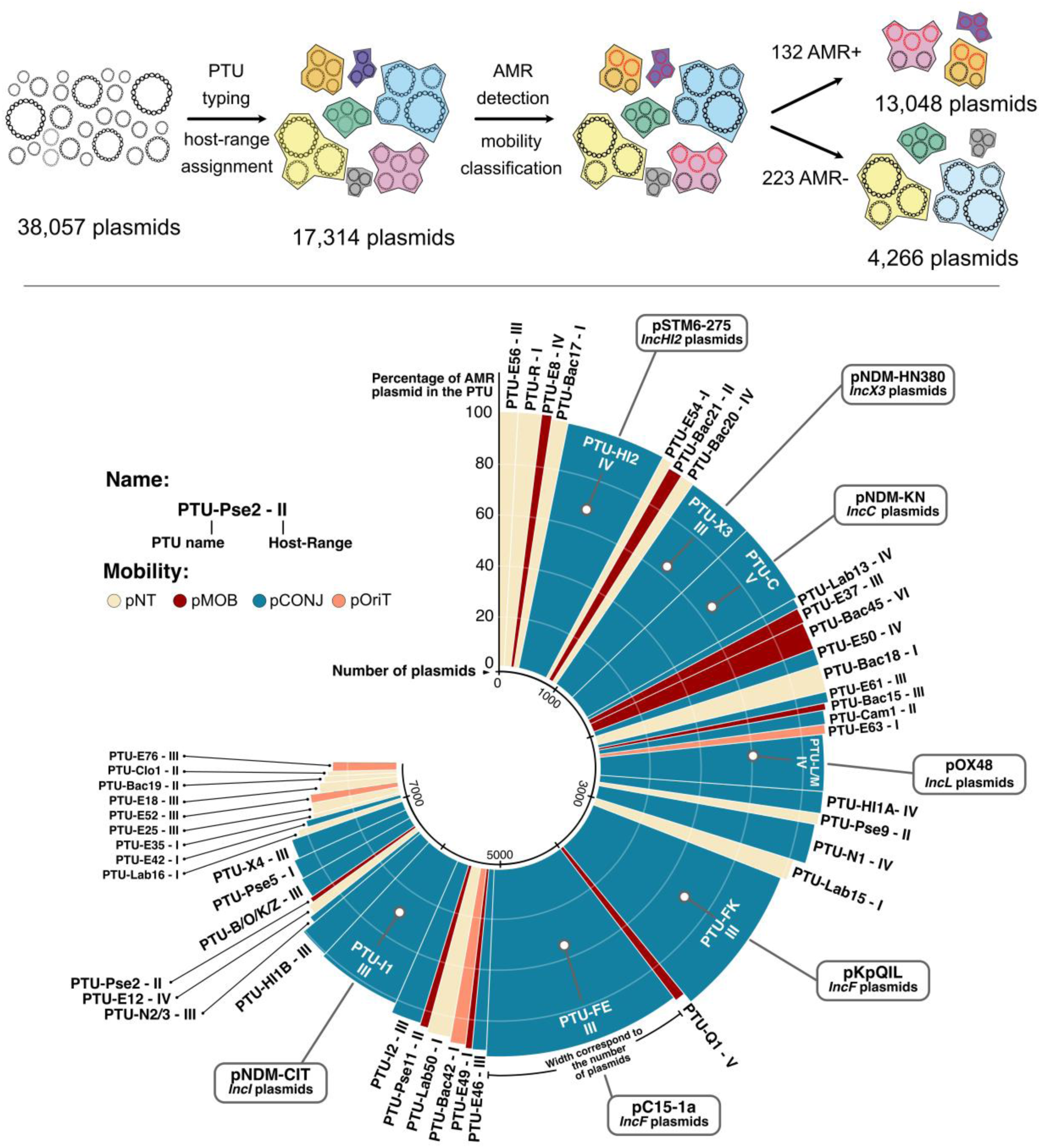
Host-Range, mobility and frequency of AMR amongst the 50 most prevalent AMR+ PTU. Top. Method to class the PTUs regarding AMR. **Bottom.** Percentage of AMR plasmids in the PTUs. The width of the bar represents the number of plasmids in the PTU and its color indicates the most frequent mobility in the PTU. The name of the PTU is written at the top of the bar and roman numbers show the host-range of the PTU. For the most abundant PTUs, representative plasmids and frequently associated incompatibility groups are shown in bubbles. pNT: non-transmissible plasmid; pOriT: plasmid carrying an origin of transfer; pMOB: mobilizable plasmid; pCONJ: conjugative plasmid.

### Resistance plasmids and their PTUs are mobile and have broad host ranges

To test the hypothesis that plasmids encoding antibiotic resistance tend to have broad host ranges, we compared the PTUs containing such plasmids with the others. For these analyses we used a previously published classification of host-ranges of PTUs [54]. In that study, host ranges were classed in six groups from I (narrowest) and VI (broadest). To have enough sample size for statistical analyses, we put together the classes V and VI. We found that the frequency of PTUs encoding AMR was higher for the broadest host ranges (Nominal logistic regression P<0.0001, Figure 3A, Table S3). As we feared that the high number of replicons in certain PTUs could inflate the PTU host range, we made a complementary analysis where we added the number of plasmids in each PTU in the statistical model. The association between PTU encoding AMR and broad host range remained significant (Nominal logistic regression P<0.0001, Table S3), and is thus not an artifact caused by the different number of plasmids per PTU.

**Figure 3:**
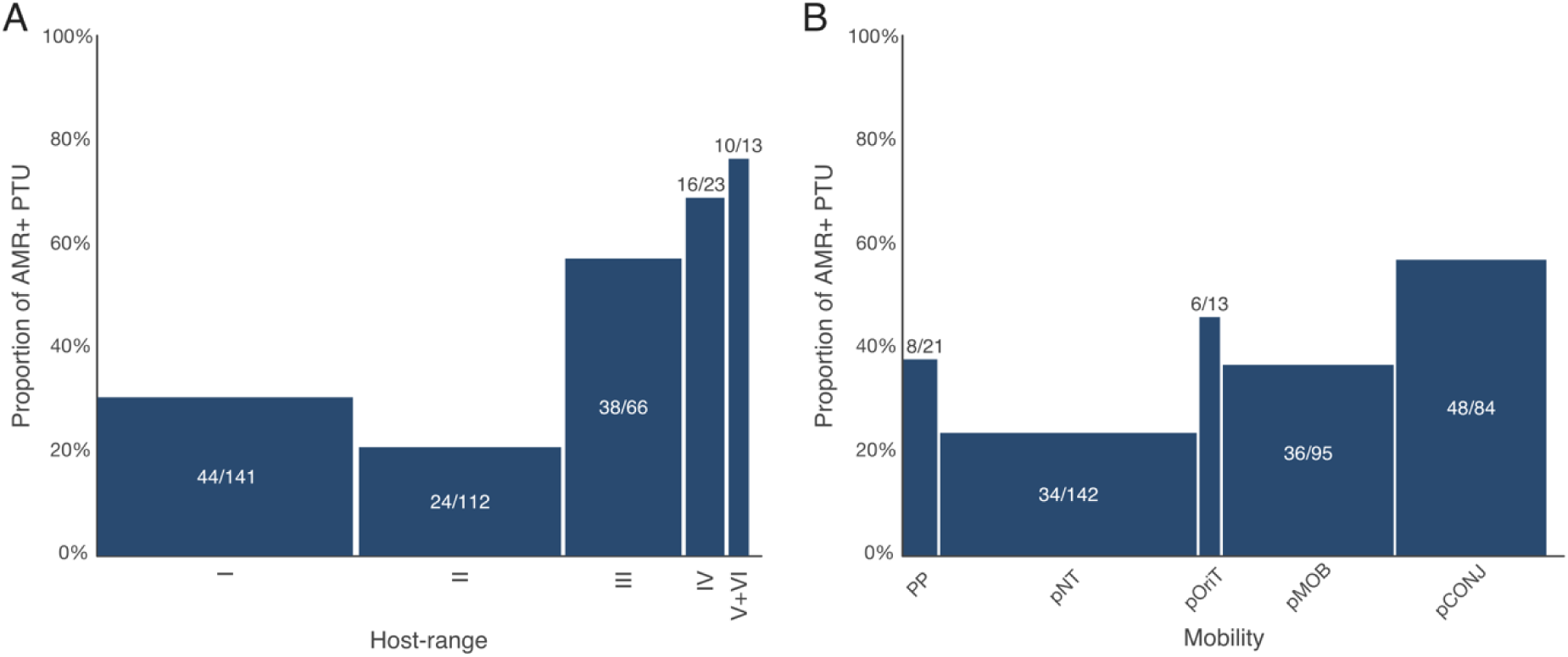
Proportion of PTUs with plasmids encoding AMR genes in function of host-range (A, I-narrowest, VI-broadest) and mobility (B). The numbers over the bars correspond to the proportion of PTUs with plasmids encoding at least one antibiotic resistance gene (AMR+). PP: Phage-plasmid; pNT: non-transmissible plasmid; pOriT: plasmid carrying an origin of transfer; pMOB: mobilizable plasmid; pCONJ: conjugative plasmid.

To study the mobility of plasmids in light of the carriage of AMR, we classified the former as: phage-plasmids (P-P), conjugative plasmids (pCONJ), mobilizable plasmids (encoding a relaxase) (pMOB), plasmids with an origin of transfer for conjugation (pOriT), and putatively non-transmissible plasmids (pNT, see Methods). We then identified the most frequent mobility type for each PTU and assigned it to the group. In 97% of the cases the major category accounted for more than 50% of the plasmids of the group and often much more (Figure S4). For instance, in 232 out of the 355 PTU (65%), more than 90% of plasmids had the same classification in terms of mobility. Our analysis shows that PTUs with a majority of conjugative plasmids are those with the highest frequency of plasmids carrying antibiotic resistance genes, followed by diverse types of mobilizable plasmids and phage-plasmids. The plasmids putatively non-transmissible had the lowest frequency of antibiotic resistance (Figure 3B, Table S4, P < 0.0001). Taken together these results suggest that plasmids encoding antibiotic resistance tend to be mobile and have broader host-ranges than the others.

### Antibiotic resistance plasmids recombine more

Plasmids may evolve rapidly by exchanging genes and alleles with other plasmids or chromosomes. Homologous recombination is usually regarded as a mechanism of allelic variation, which can speed up adaptation. But recombination involving multiple genes may also result in gene acquisition or loss. The numerous plasmids in some PTUs allow to identify homologous recombination in their core genes. We found 270 PTUs having between 1 and 381 gene families present in more than 80% of the plasmids (defined as core genes of the PTU, Figure S5). As expected, the number of core genes in a PTU correlates with the mean size of the plasmids in the PTU (Spearman correlation: 0.7033, P< 0.0001). Importantly, there is no significant difference in the number of core genes between PTUs with or without AMR genes (Wilcoxon test, P= 0.482). While this may seem surprising given that plasmids encoding AMR are larger (see above), many of the smallest plasmids could not be placed in PTUs and the number of core genes is shaped by the size of the element and its rate of variation of gene repertoires (see below). Hence, whatever differences one may find between these two groups of PTUs, they are not caused by differences in the number of their core genes.

We analyzed the rates of recombination using two variants of a phylogenetic method. In method I we compared the evolutionary history of each core gene in the PTU with that of the concatenate of the core genes of the PTU (henceforth called the consensus phylogeny). This was done using the normalized Robinson-Foulds (nRF) distances which measure the similarity between two phylogenetic trees (Figure 4). This analysis involved the analysis of 11,294 gene trees from the 250 PTUs with more than one core gene. 17 PTUs had congruent trees, i.e the evolutionary histories of individual core genes were similar from the consensus, suggesting little or no recombination (mean nRF=0, Table S2). Most of the others showed evidence of extensive recombination. The average incongruence was high (median nRF of 0.505), i.e., the evolutionary histories of the individual core genes were quite different from the consensus. If there are high recombination rates in plasmids, then the phylogeny of the concatenate of the core genes may not be reliable. Hence, we developed a complementary approach (method II, Figure 4) where we compared the average nRF between every pair of gene families in the PTU. This analysis does not require the use of a consensus trees and results in similar conclusions (median nRF of 0.59, Table S5), confirming that recombination is frequent in plasmid core genes.

**Figure 4:**
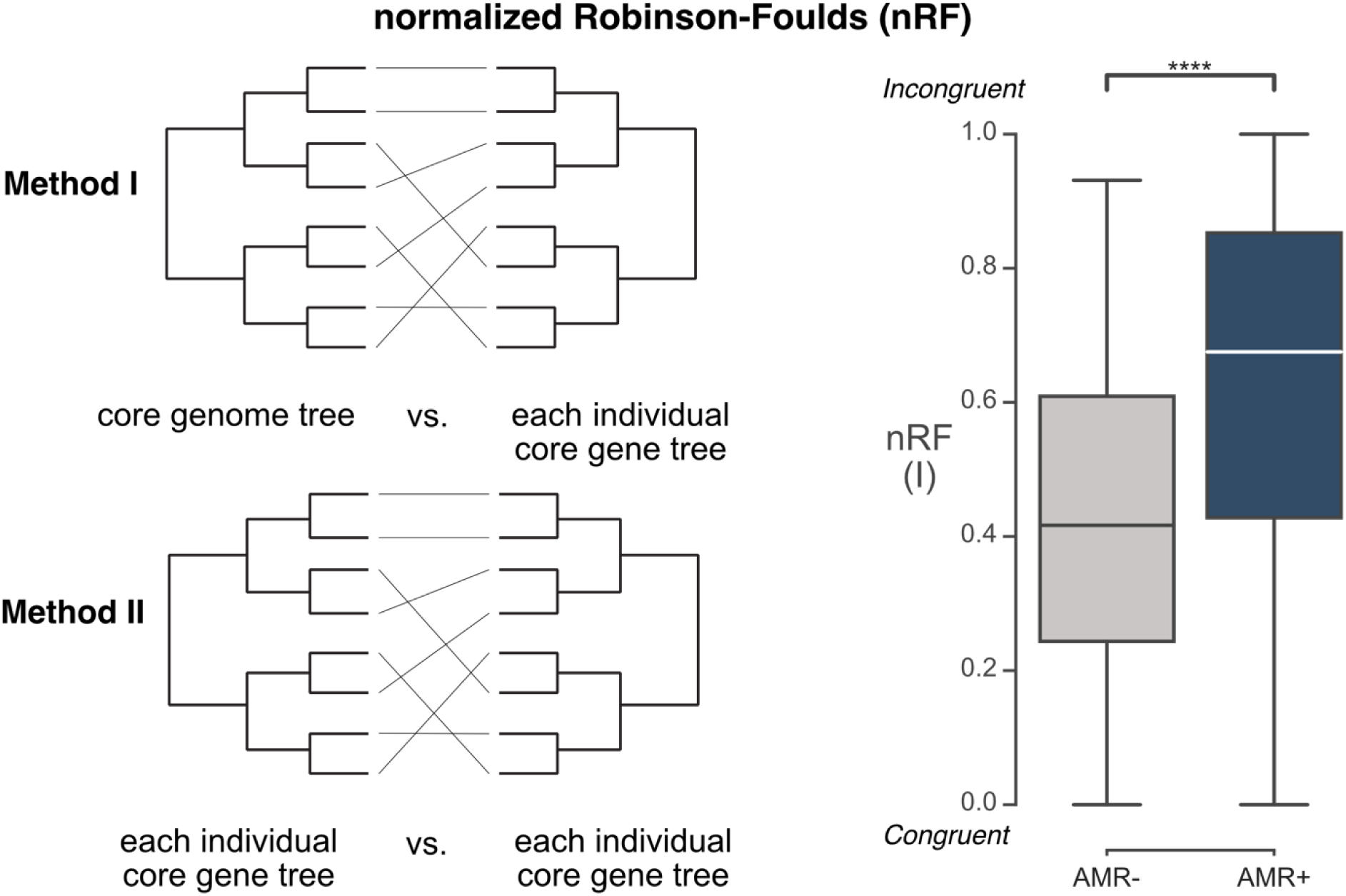
Phylogenetic incongruence in PTUs encoding (AMR+) or lacking (AMR-) antibiotic resistance. The average normalized Robinson-Foulds (nRF) for a PTU was computed between the PTU core genome tree and the tree of each core gene or between every pair of trees of the core genes (called Method I in the text). Boxplots are represented as in Figure 1. Statistical test (Mann-Whitney U): **** P ≤ 0.0001.

To test the idea that plasmids encoding AMR recombine at higher rates than the others, we compared the average incongruence (obtained using method I) between the PTUs with or without AMR genes. In agreement with our hypothesis, incongruences were higher in PTUs encoding AMR (Figure 4, Nominal logistic regression, P<0.0001, Table S6). A complementary statistical model including the number of core genes and the size of the plasmids as covariates confirmed the significance of the positive association between recombination (measured by nRF) and the presence of antibiotic resistance in PTUs (P<0.0001, same test, Table S6). This result is not caused by different depths in the phylogenetic trees of the two groups of PTUs, since they are not significantly different in this respect (Figure S6). A similar type of association between nRF and AMR was found with the method II (P<0.0001, same test, Table S5). Overall, these results show that PTUs with antibiotic resistance plasmids engage in allelic exchanges at higher rates than the others.

### Gene turnover is higher in plasmid families encoding AMR

If plasmids with antibiotic resistance genes are more plastic, then their PTUs might have a higher diversity of gene families, i.e. larger pangenomes. To test this hypothesis, we computed the rarefaction curves of the pangenomes for PTUs with more than 30 plasmids and fitted them with the Heaps’ law (see Methods). We then analyzed the β parameter that varies between 0 (close, invariant genome) and 1 (open, highly variable genome) [49](Figure 5A). The pangenomes of PTUs are open (median β is 0.526), and those with antibiotic resistance genes are more open than the others (β higher by 17.5%, Figure 5A, P=0.0007, Nominal logistic regression, Table S7). As the number of plasmids in the PTU and the size of the plasmids could have an impact on the saturation curves, we made an additional model including both parameters. This complementary analysis confirmed a significantly higher average value of β in the PTUs containing AMR-genes (P=0.004, same test, Table S7). These results indicate that plasmids with antibiotic resistance genes have a more open pangenome.

**Figure 5:**
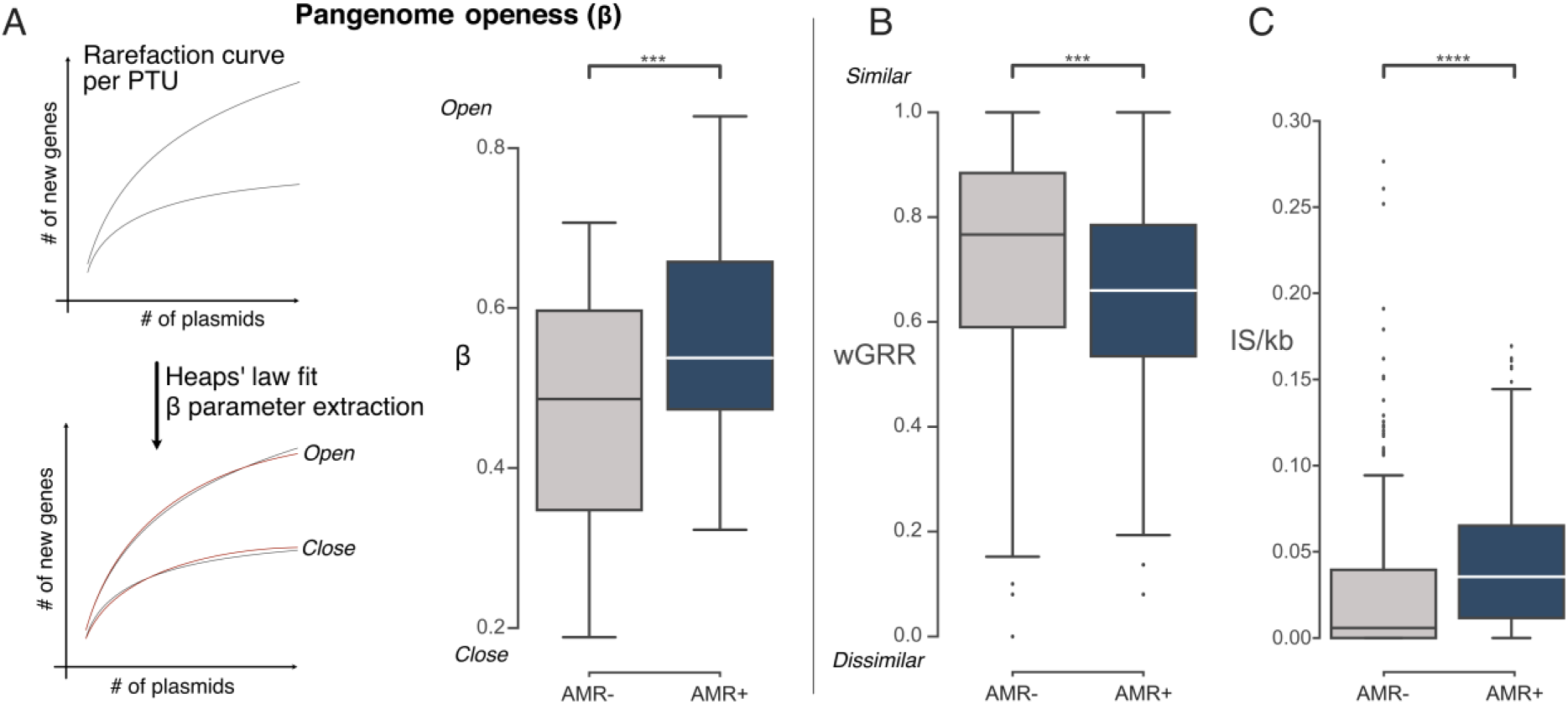
Distribution of the mean beta (A, with explanatory schema) and wGRR values (B) and IS density (C) of PTUs encoding (AMR+) or lacking (AMR-) antibiotic resistance genes. The beta parameters of the heaps’ law were extracted from the fit of the rarefaction curve for each PTU. Boxplots are represented as in Figure 1. Statistical tests (Mann-Whitney U): *** P ≤ 0.001; **** P ≤ 0.0001.

If the pangenomes of PTUs with antibiotic resistance plasmids are more open, then these plasmids should be more diverse in terms of accessory genes. To test this hypothesis, we compared the divergence in gene repertoires between all pairs of plasmids of the same PTU with the weighted Gene Repertoire Relatedness index (wGRR, see Methods). The wGRR varies between 0 and 1. A wGRR close to 1 means that both plasmids are very similar (all genes in a plasmid have a very similar BBH in the other) and a wGRR o 0 means that plasmids lack homologs. As expected, PTUs with antibiotic resistance genes had a significantly smaller wGRR than the others (Figure 5B). When we fitted a nominal logistic model between the AMR status of a PTU and the mean wGRR, smaller values of wGRR were associated with PTUs with antibiotic resistance genes (Nominal logistic regression, P=0.0006), even controlling for the number of core genes and the number of plasmids in the PTU (Table S8). Hence, despite the high similarity of core genes within PTUs, their plasmids have highly variable gene repertoires. PTUs encoding AMR are even more variable than the others.

Acquisition and exchange of AMR genes in plasmids is often driven by integrons and/or transposons (see Introduction). Indeed, we found that PTUs encoding AMR harbor higher densities of transposable elements and are the only ones with integrons (Figure 5C/Figure S7). These results suggest that, while having similar numbers of core genes, the PTUs with antibiotic resistance genes are enriched in genetic elements that increase the rates of gain and loss (turnover) of accessory genes.

PTUs with antibiotic resistance genes have higher recombination rates in core genes, higher gene repertoire diversity, and higher gene turnover. Since these factors are not independent, we assessed their joint effect. All three variables are significantly and positively associated with PTUs with antibiotic resistance genes (P< 0.0001, Nominal logistic regression, Table S9), suggesting that all these factors favor the acquisition of antibiotic resistance genes. We then compared the Wald χ_²_ and the odds ratios of these factors to determine which had the strongest effect. The openness of the pangenome had the strongest effect (beta_χ²_ = 11.58, Odds ratio: 7995.991), followed by the gene repertoire diversity (wGRR_χ²_ = 8.79, Odds ratio: 0.003466), and the core genome recombination rate (nRF_χ²_ = 5.38, Odds ratio: 70.631). This suggests that while all these factors contribute to the acquisition of AMR genes by plasmids, the capacity to quickly acquire new genes (resulting in a more open pangenome and higher wGRR) is particularly important.

### Evolvability and mobility are intrinsic to the AMR PTUs

The previous results showed that PTUs with antibiotic resistance genes are more plastic than the others. This can be interpreted in two ways. 1) Plasmids that are intrinsically more plastic and mobile are more likely to acquire resistance genes and spread them. 2) Resistance itself favors rapid genetic diversification and mobility. The second hypothesis is consistent with the association of resistance genes with recombinases (IS, integrons), although the way resistance could favor plasmid mobility is less obvious. To better understand the relations of causality, we kept the classification of AMR-association of PTUs (AMR+ and AMR-) but excluded the resistance plasmids from the analysis (Figure 6). AMR+ PTUs contained more conjugative elements and had broader host-ranges than the others (Table S10, P < 0.0001). They also had higher recombination rates, higher mean nRF values (P = 0.0170, nominal logistic regression, Table S10, Figure 6A), lower wGRR, and more open pangenomes (P_beta_ = 0.0027, P_wGRR_ = 0.0006, same test, Table S10, Figure 6B/C). The density of transposable elements was higher in these PTUs (P =0.01025, Figure 6D). Hence, the rapid evolution of antibiotic resistance plasmids seems intrinsic to the PTU and is not just a consequence of the presence of resistance genes in some plasmids of the PTU. This suggests a causality: more plastic and mobile PTUs are more likely to acquire antibiotic resistance genes and spread them among human-related bacteria.

**Figure 6:**
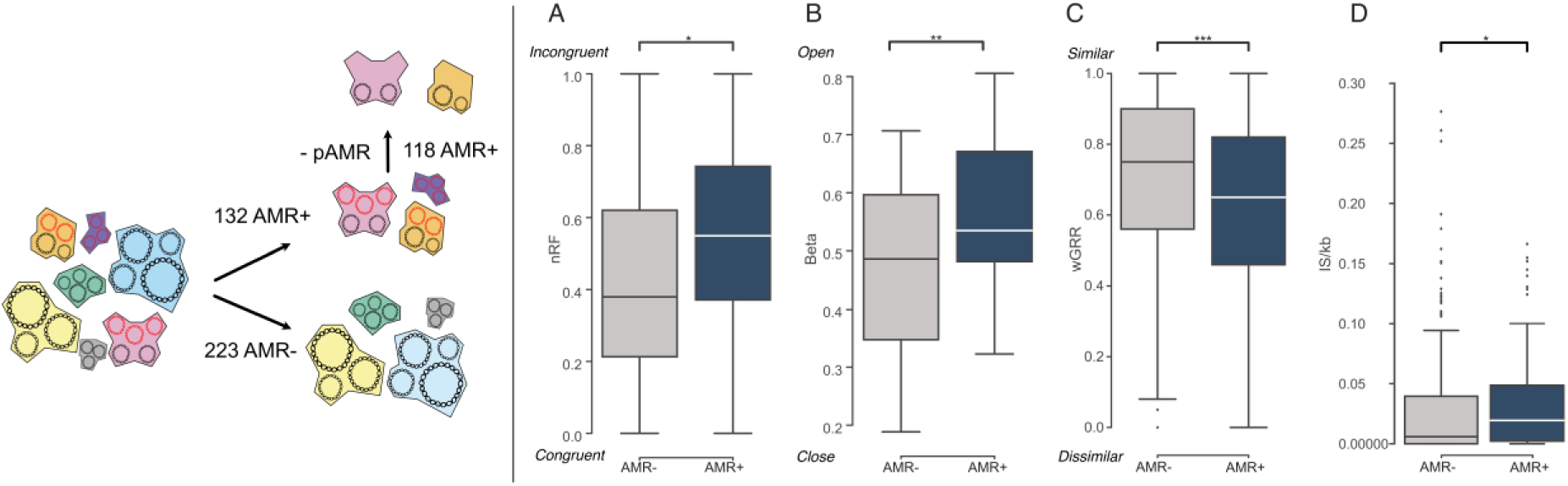
Analysis of PTUs (as in Figure 2), but where the PTU AMR+ were purged from antibiotic resistance plasmids. **Left panel:** Schema describing dataset filtering and number of PTUs in the final dataset**. Right panels**: Distribution of the values of mean nRF (A), beta (B), mean wGRR (C) and IS density (D) between AMR- and AMR+ PTUs (excluding the antibiotic resistance plasmids). Boxplots are presented as in Figure 1. Statistical significance (Mann-Whitney U): * P ≤ 0.05; ** P ≤ 0.01; ***: p ≤ 0.001.

## Discussion

Here, we focused on how a trait (antibiotic resistance) can be acquired by horizontal gene transfer when bacteria need to tackle an unprecedented challenge. Antibiotic resistance in nosocomials differs from traits like virulence, competition, or phage immunity by its extreme novelty. This is why we focused on known antibiotic resistance genes of relevance in the clinic, as they are recent in these bacteria. Environmental bacteria with other antibiotic resistance genes may endure different dynamics because we cannot ensure such trait is recent in their habitat. It is not surprising that conjugative plasmids have the highest frequency of such AMR genes in our dataset, since they are mobile, tend to have broad host range, can transfer large amounts of genetic information per event, and tolerate large integrations of genetic material [55]. Plasmids that are not autonomously conjugative but can be mobilized by conjugation also account for a large fraction of transfer of resistance genes, and some of them have amongst the broadest known host ranges [56]. Future work will be needed to include the substantial number of plasmids, mostly small, that could not be classed in PTUs. Overall, our results are in line with the hypothesis that broad-host range conjugation has a key role in driving the initial stages of transfer of novel traits between distantly related bacteria.

By focusing on homogeneous groups of closely related plasmids (PTUs) we could analyze these plasmids in a phylogenetic context to analyze their rates of change and of recombination. Unfortunately, linking all the PTUs by a phylogeny is impossible due to the lack of common genes and we treated the PTUs as effectively independent units. These analyses confirmed that families of plasmids encoding antibiotic resistance have larger pangenomes and faster-changing gene repertoires than the others. We cannot exclude the possibility that the bias in genome databases towards closely related antibiotic resistant bacteria may result in larger numbers of plasmids in PTUs encoding AMR. This should make our analysis conservative, since more closely related plasmids are more likely to reveal higher average values of gene repertoire relatedness (wGRR) and we found the opposite trend. It could also be argued that antibiotic resistance plasmids inflate estimates of the PTUs rate of diversification of gene repertoires because we know these genes were acquired recently. Yet, the analysis of PTUs after removing antibiotic resistance plasmids corroborated our hypothesis that evolvability favors AMR acquisition.

The acquisition of AMR is likely facilitated by the abundance of transposable elements and integrons in plasmids, as previously described for resistance plasmids [22] and here further extended to their PTUs (even when removing AMR plasmids from the analyses). The reasons for the higher frequency of transposable elements in some plasmids are poorly understood. It may be related with the availability of more neutral integration sites in the larger conjugative plasmids or with preferential transposition during plasmid conjugation [57]. It is important to note that transposable elements may also facilitate the transfer of resistance from plasmids to chromosomes and this might stabilize resistance in the genome at a later stage [58]. It is thus possible that the same mobile genetic elements that facilitated the acquisition of antibiotic resistance by highly mobile and plastic plasmids will subsequently have a key role in the transfer of these genes to more stable regions of the bacterial genome.

The relevance of homologous recombination in the acquisition of AMR by plasmids has rarely been studied, even though it was previously observed that homologous recombination was frequent in core genes of IncP-1 plasmids [59]. Actually, the extent of recombination, measured by phylogenetic incongruence, was very different between PTUs, and was particularly high in PTUs encoding AMR. It’s important to note that homologous recombination may result in the acquisition of intervening genes when recombination tracts are long [60]. Hence, homologous recombination may be part of the mechanisms allowing the plasmids to acquire resistance genes. The abundance of transposable elements in AMR plasmids may also increase the chances of DNA breaks that stimulate repair by homologous recombination [61, 62], and thereby stimulate allelic exchanges. Plasmid copy number, when high, may also result in higher frequency of genetic exchanges [63]. Finally, the mobility of antibiotic resistance plasmids may increase the likelihood of their occasional co-occurrence with other plasmids, which may result in more frequent and varied genetic exchanges [21]. In the future, it would be interesting to compare these results with those of plasmids carrying other types of traits like virulence or defense to understand if given time antibiotic resistance genes will tend to move towards plasmids with other characteristics such as low cost and high stability.

Phages can disperse freely in the environment and might be better at transferring novel genes between biomes. But temperate phages tend to have narrow host ranges caused by their dependency on specific receptor binding proteins [64]. These phages are also less tolerant to gene acquisition than conjugative elements because increase in genome size rapidly results in defaults in DNA packaging in the viral particle [65]. Our analyses highlighted that variability in gene repertoire changes seems to be the most important trait for the acquisition of AMR genes. This may explain why integrative phages tend to lack such genes [66] since they are very constrained in size. In contrast, phage-plasmids play a role in the spread of antibiotic resistance possibly because they may accept novel genes more easily (they are larger), they have more transposable elements and integrons, and they exchange genes with plasmids at larger rates [19]. Interestingly, P1-like phage-plasmids, which are particularly large and of broad host range [67, 68], are among the ones having most resistances (Figure S3). This suggests that host range and evolvability also facilitate the transfer of novel traits by phages. Of note, while conjugation seems to have a key role in the initial transfer of traits across biomes and distantly related bacteria [22], once the antibiotic resistance genes are acquired by nosocomial bacteria, phage-mediated transduction and natural transformation may further spread them between closely related bacteria [69].

PTUs encoding antibiotic resistance are associated with conjugation, broad host range, and several forms of genetic diversification, even when we removed antibiotic resistance plasmids from the dataset. This suggests that these are intrinsic characteristics of the plasmid families that acquire AMR, and not the result of antibiotic resistance acquisition. That some plasmid families are more prone to the transfer of adaptive genes from distant bacteria in different biomes, implies that plasmids have differentiated roles in the processes of horizontal gene transfer. This may be of importance to study the transfer of traits tackling other unprecedented challenges, such as response to pollution or climate change.

## Acknowledgements

We thank Manuel Ares-Arroyo and Marie Touchon for discussions and comments on the manuscript. We thank Eugen Pfeifer for providing the phage-plasmid classification. This work used the computational and storage services (TARS cluster) provided by the IT department at Institut Pasteur, Paris. Work was supported by the Laboratoire d’Excellence IBEID Integrative Biology of Emerging Infectious Diseases [ANR-10-LABX-62-IBEID], Equipe FRM (Fondation pour la Recherche Médicale) [EQU201903007835], and the INCEPTION project [PIA/ANR-16-CONV-0005].

## Bibliography

1. Arnold, B.J., I.T. Huang, and W.P. Hanage, Horizontal gene transfer and adaptive evolution in bacteria. Nat Rev Microbiol, 2022. 20(4): p. 206–218.

2. Brockhurst, M.A., et al., The Ecology and Evolution of Pangenomes. Current Biology, 2019. 29(20): p. R1094–R1103.

3. Cabezon, E., et al., Towards an integrated model of bacterial conjugation. FEMS Microbiol Rev, 2015. 39(1): p. 81–95.

4. Rankin, D.J., E.P.C. Rocha, and S.P. Brown, What traits are carried on mobile genetic elements, and why? Heredity, 2011. 104(1): p. 1–10.

5. Hedrick, P.W., Balancing selection. Curr Biol, 2007. 17(7): p. R230–1.

6. San Millan, A., et al., Multicopy plasmids potentiate the evolution of antibiotic resistance in bacteria. Nature ecology & evolution, 2016. 1(1): p. 1–8.

7. Ilhan, J., et al., Segregational drift and the interplay between plasmid copy number and evolvability. Molecular biology and evolution, 2019. 36(3): p. 472–486.

8. Nicolaou, K.C. and S. Rigol, A brief history of antibiotics and select advances in their synthesis. The Journal of Antibiotics, 2018. 71(2): p. 153–184.

9. Larsson, D.G.J. and C.F. Flach, Antibiotic resistance in the environment. Nature Reviews Microbiology, 2022. 20(5): p. 257–269.

10. Castaneda-Barba, S., E.M. Top, and T. Stalder, Plasmids, a molecular cornerstone of antimicrobial resistance in the One Health era. Nat Rev Microbiol, 2024. 22(1): p. 18–32.

11. Hughes, V.M. and N. Datta, Conjugative plasmids in bacteria of the ‘pre-antibiotic’ era. Nature, 1983. 302(5910): p. 725–726.

12. Bean, D.C., D.M. Livermore, and L.M. Hall, Plasmids imparting sulfonamide resistance in Escherichia coli: implications for persistence. Antimicrob Agents Chemother, 2009. 53(3): p. 1088–93.

13. Rozwandowicz, M., et al., Plasmids carrying antimicrobial resistance genes in Enterobacteriaceae. J Antimicrob Chemother, 2018. 73(5): p. 1121–1137.

14. Wang, R., et al., The global distribution and spread of the mobilized colistin resistance gene mcr-1. Nat Commun, 2018. 9(1): p. 1179.

15. León-Sampedro, R., et al., Pervasive transmission of a carbapenem resistance plasmid in the gut microbiota of hospitalized patients. Nature Microbiology, 2021. 6(5): p. 606–616.

16. Kondratyeva, K., M. Salmon-Divon, and S. Navon-Venezia, Meta-analysis of Pandemic ST131 Plasmidome Proves Restricted Plasmid-clade Associations. Scientific Reports, 2020. 10(1).

17. Deleo, F.R., et al., Molecular dissection of the evolution of carbapenem-resistant multilocus sequence type 258 Klebsiella pneumoniae. Proc Natl Acad Sci U S A, 2014. 111(13): p. 4988–93.

18. Botelho, J. and H. Schulenburg, The Role of Integrative and Conjugative Elements in Antibiotic Resistance Evolution. Trends in Microbiology, 2021. 29(1): p. 8–18.

19. Pfeifer, E., R.A. Bonnin, and E.P.C. Rocha, Phage-Plasmids Spread Antibiotic Resistance Genes through Infection and Lysogenic Conversion. Mbio, 2022. 13(5).

20. Amos, G.C.A., et al., The widespread dissemination of integrons throughout bacterial communities in a riverine system. ISME J, 2018. 12(3): p. 681–691.

21. Wang, X., et al., Inter-plasmid transfer of antibiotic resistance genes accelerates antibiotic resistance in bacterial pathogens. The ISME Journal, 2024. 18(1): p. wrad032.

22. Che, Y., et al., Conjugative plasmids interact with insertion sequences to shape the horizontal transfer of antimicrobial resistance genes. Proc Natl Acad Sci U S A, 2021. 118(6).

23. Deng, Y., et al., Resistance integrons: class 1, 2 and 3 integrons. Ann Clin Microbiol Antimicrob, 2015. 14: p. 45.

24. Wang, Y. and T. Dagan, The evolution of antibiotic resistance islands occurs within the framework of plasmid lineages. Nature Communications, 2024. 15(1): p. 4555.

25. Wang, Y.Q., et al., Gene sharing among plasmids and chromosomes reveals barriers for antibiotic resistance gene transfer. Philosophical Transactions of the Royal Society B-Biological Sciences, 2022. 377(1842).

26. Sekizuka, T., et al., Complete sequencing of the bla(NDM-1)-positive IncA/C plasmid from Escherichia coli ST38 isolate suggests a possible origin from plant pathogens. PLoS One, 2011. 6(9): p. e25334.

27. Lipworth, S., et al., The plasmidome associated with Gram-negative bloodstream infections: A large-scale observational study using complete plasmid assemblies. Nature Communications, 2024. 15(1): p. 1612.

28. Anda, M., et al., Bacteria can maintain rRNA operons solely on plasmids for hundreds of millions of years. Nature Communications, 2023. 14(1): p. 7232.

29. Hutchings, M.I., A.W. Truman, and B. Wilkinson, Antibiotics: past, present and future. Curr Opin Microbiol, 2019. 51: p. 72–80.

30. O’Leary, N.A., et al., Reference sequence (RefSeq) database at NCBI: current status, taxonomic expansion, and functional annotation. Nucleic Acids Res, 2016. 44(D1): p. D733–45.

31. Hyatt, D., et al., Prodigal: prokaryotic gene recognition and translation initiation site identification. BMC Bioinformatics, 2010. 11: p. 119.

32. Néron, B., et al., MacSyFinder v2: Improved modelling and search engine to identify molecular systems in genomes. Peer Community Journal, 2023. 3.

33. Coluzzi, C., et al., Evolution of Plasmid Mobility: Origin and Fate of Conjugative and Nonconjugative Plasmids. Molecular Biology and Evolution, 2022. 39(6): p. msac115.

34. Camacho, C., et al., BLAST+: architecture and applications. BMC Bioinformatics, 2009. 10: p. 421.

35. Ares-Arroyo, M., C. Coluzzi, and E.P.C. Rocha, Origins of transfer establish networks of functional dependencies for plasmid transfer by conjugation. Nucleic Acids Research, 2023. 51(7): p. 3001–3016.

36. Pfeifer, E., et al., Bacteria have numerous distinctive groups of phage–plasmids with conserved phage and variable plasmid gene repertoires. Nucleic Acids Research, 2021. 49(5): p. 2655–2673.

37. Feldgarden, M., et al., AMRFinderPlus and the Reference Gene Catalog facilitate examination of the genomic links among antimicrobial resistance, stress response, and virulence. Sci Rep, 2021. 11(1): p. 12728.

38. Xie, Z.Q. and H.X. Tang, ISEScan: automated identification of insertion sequence elements in prokaryotic genomes. Bioinformatics, 2017. 33(21): p. 3340–3347.

39. Neron, B., et al., IntegronFinder 2.0: Identification and Analysis of Integrons across Bacteria, with a Focus on Antibiotic Resistance in Klebsiella. Microorganisms, 2022. 10(4).

40. Redondo-Salvo, S., et al., COPLA, a taxonomic classifier of plasmids. BMC Bioinformatics, 2021. 22(1): p. 390.

41. Perrin, A. and E.P.C. Rocha, PanACoTA: a modular tool for massive microbial comparative genomics. NAR Genom Bioinform, 2021. 3(1): p. lqaa106.

42. Steinegger, M. and J. Söding, MMseqs2 enables sensitive protein sequence searching for the analysis of massive data sets. Nature biotechnology, 2017. 35(11): p. 1026.

43. Katoh, K. and D.M. Standley, MAFFT: iterative refinement and additional methods. Methods Mol Biol, 2014. 1079: p. 131–46.

44. Nguyen, L.T., et al., IQ-TREE: a fast and effective stochastic algorithm for estimating maximum-likelihood phylogenies. Mol Biol Evol, 2015. 32(1): p. 268–74.

45. Kalyaanamoorthy, S., et al., ModelFinder: fast model selection for accurate phylogenetic estimates. Nature methods, 2017. 14(6): p. 587.

46. Filipski, A., et al., Prospects for building large timetrees using molecular data with incomplete gene coverage among species. Molecular Biology and Evolution, 2014. 31(9): p. 2542–2550.

47. Huerta-Cepas, J., F. Serra, and P. Bork, ETE 3: Reconstruction, Analysis, and Visualization of Phylogenomic Data. Mol Biol Evol, 2016. 33(6): p. 1635–8.

48. Newton, A., Forest Ecology and Conservation : A Handbook of Techniques. Techniques in Ecology & Conservation. 2007, Oxford: Oxford University Press.

49. Tettelin, H., et al., Comparative genomics: the bacterial pan-genome. Curr Opin Microbiol, 2008. 11(5): p. 472–7.

50. Virtanen, P., et al., SciPy 1.0: fundamental algorithms for scientific computing in Python. Nat Methods, 2020. 17(3): p. 261–272.

51. Robinson, D.F. and L.R. Foulds, Comparison of phylogenetic trees. Mathematical Biosciences, 1981. 53: p. 131–147.

52. Didelot, X. and D. Falush, Inference of bacterial microevolution using multilocus sequence data. Genetics, 2007. 175: p. 1251–66.

53. Cury, J., et al., Host Range and Genetic Plasticity Explain the Coexistence of Integrative and Extrachromosomal Mobile Genetic Elements. Mol Biol Evol, 2018. 35(11): p. 2230– 2239.

54. Redondo-Salvo, S., et al., Pathways for horizontal gene transfer in bacteria revealed by a global map of their plasmids. Nature communications, 2020. 11(1): p. 1–13.

55. Forster, S.C., et al., Strain-level characterization of broad host range mobile genetic elements transferring antibiotic resistance from the human microbiome. Nature Communications, 2022. 13(1): p. 1445.

56. Meyer, R., Replication and conjugative mobilization of broad host-range IncQ plasmids. Plasmid, 2009. 62(2): p. 57–70.

57. Ton-Hoang, B., et al., Single-Stranded DNA Transposition Is Coupled to Host Replication. Cell, 2010. 142(3): p. 398–408.

58. Kadibalban, A.S., G. Landan, and T. Dagan, The extent and characteristics of DNA transfer between plasmids and chromosomes. Curr Biol, 2024.

59. Norberg, P., et al., The IncP-1 plasmid backbone adapts to different host bacterial species and evolves through homologous recombination. Nat Commun, 2011. 2: p. 268.

60. Doroghazi, J.R. and D.H. Buckley, Widespread homologous recombination within and between species. ISME J, 2010. 4(9): p. 1136–1143.

61. Li, X. and W.-D. Heyer, Homologous recombination in DNA repair and DNA damage tolerance. Cell Research, 2008. 18(1): p. 99–113.

62. Mahillon, J. and M. Chandler, Insertion Sequences. Microbiol Mol Biol Rev, 1998. 62: p. 725–774.

63. Nicoloff, H., et al., Three concurrent mechanisms generate gene copy number variation and transient antibiotic heteroresistance. Nature Communications, 2024. 15(1): p. 3981.

64. de Jonge, P.A., et al., Molecular and Evolutionary Determinants of Bacteriophage Host Range. Trends in Microbiology, 2019. 27(1): p. 51–63.

65. Hatfull, G.F. and R.W. Hendrix, Bacteriophages and their genomes. Current Opinion in Virology, 2011. 1(4): p. 298–303.

66. Enault, F., et al., Phages rarely encode antibiotic resistance genes: a cautionary tale for virome analyses. ISME J, 2017. 11(1): p. 237–247.

67. Murooka, Y. and T. Harada, Expansion of the host range of coliphage P1 and gene transfer from enteric bacteria to other gram-negative bacteria. Appl Environ Microbiol, 1979. 38(4): p. 754–7.

68. Pfeifer, E. and E.P.C. Rocha, Phage-plasmids promote recombination and emergence of phages and plasmids. Nature Communications, 2024. 15(1): p. 1545.

69. Winter, M., et al., Antimicrobial resistance acquisition via natural transformation: context is everything. Current Opinion in Microbiology, 2021. 64: p. 133–138.

