## Supplementary Figures for "The spread of antibiotic resistance is driven by plasmids amongst the fastest evolving and of broadest host range"

### Supplementary material

**Table of contents**

*Tables:*

Table S1: Plasmids used in the study.

Table S2: Plasmids Taxonomic Units in our dataset.

Table S3: Nominal logistic regression analysis for the host-range.

Table S4: Nominal logistic regression analysis for the mobility.

Table S5: Nominal logistic regression analysis for normalized Robinson-Foulds distance II.

Table S6: Nominal logistic regression analysis for normalized Robinson-Foulds distance I.

Table S7: Nominal logistic regression analysis for the beta parameters.

Table S8: Nominal logistic regression analysis for wGRR distance.

Table S9: Nominal logistic regression analysis for the three variables.

Table S10: Statistical analyses for the pAMR free dataset.

*Figures:*

Figure S1: Distribution of the size of plasmids.

Figure S2: Percentage of plasmids carrying AMR genes among each PTUs.

Figure S3: Distribution of unique combinations of AMR classes per PTU.

Figure S4: Proportion of the most abundant mobility type per PTU.

Figure S5: Distribution of the number of core genes per PTU.

Figure S6: Distribution of the average tips to root distance between the core genome trees of AMR+ and AMR- PTUs.

Figure S7: Distribution of the average number of complete integrons per plasmid between AMR+ and AMR- PTUs.

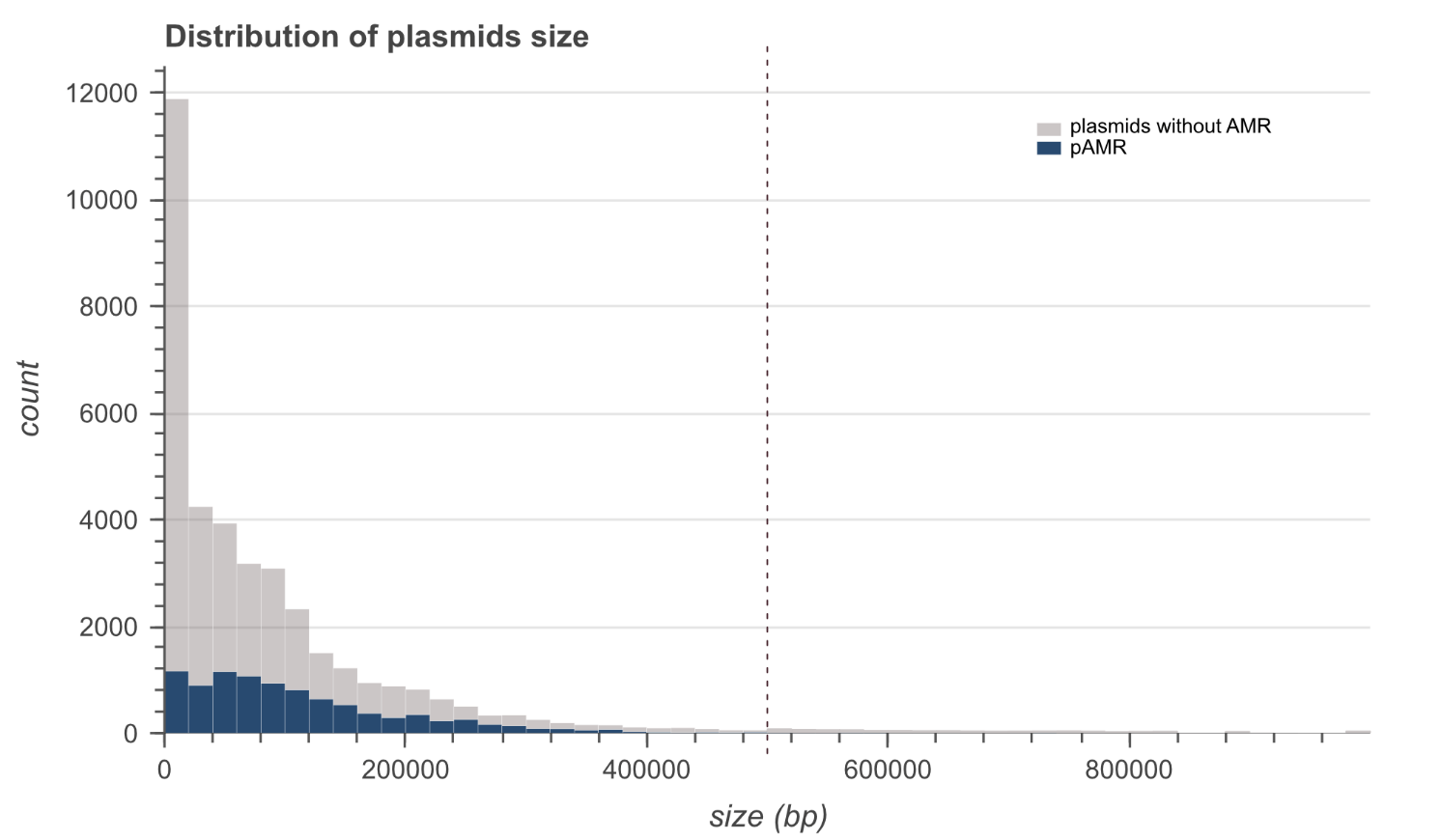

**Figure S1: Distribution of the size of plasmids encoding AMR (pAMR) or not.** Plasmids carrying AMR are shown in dark blue, while the others are in grey. The dashed line represents the 500,000 bp size limit we used to exclude the 996 large plasmids and avoid including supplementary chromosomes in our plasmid dataset.

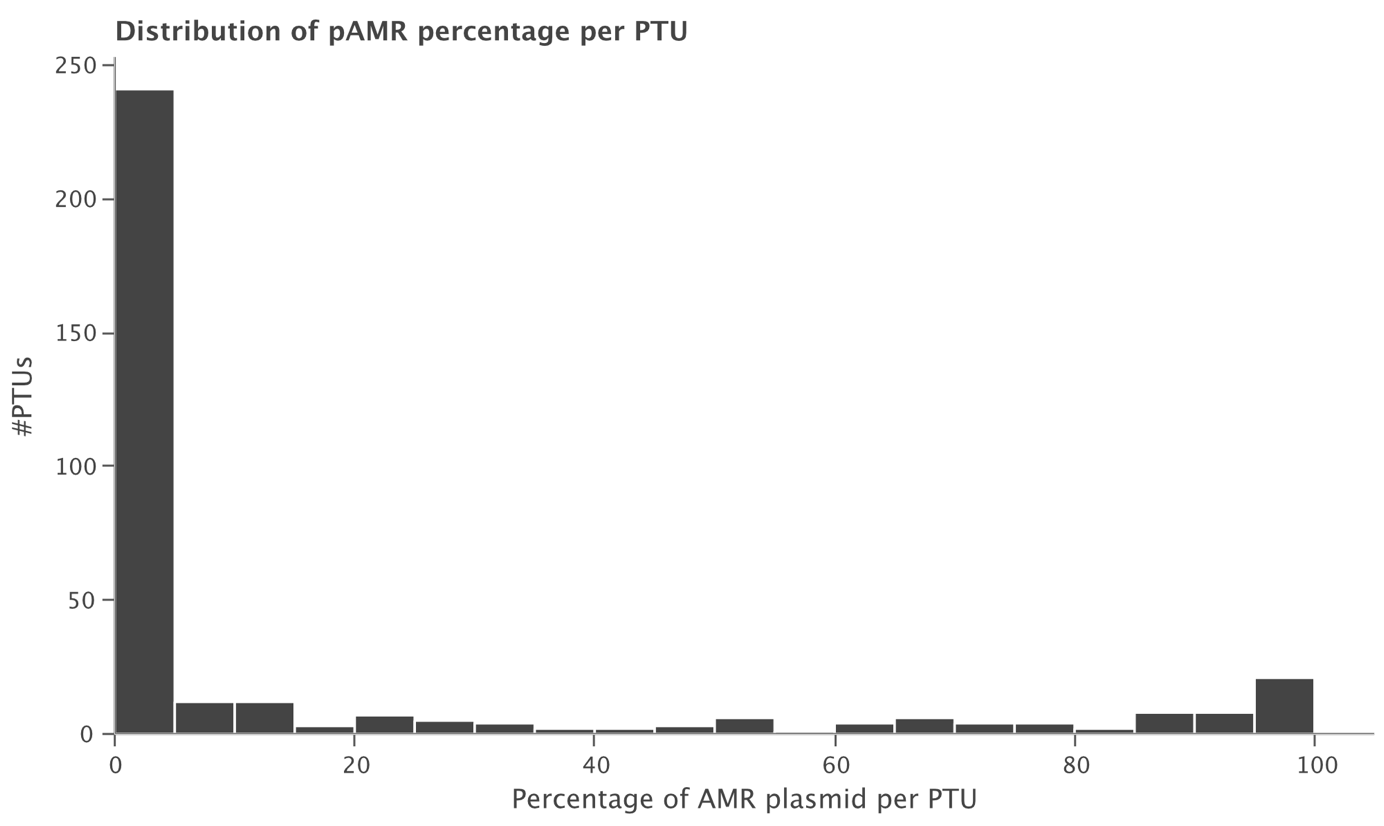
**Figure S2: Percentage of plasmids carrying AMR genes among each PTU.** In total, 223 PTUs lack AMR genes and 132 PTU contain at least one.

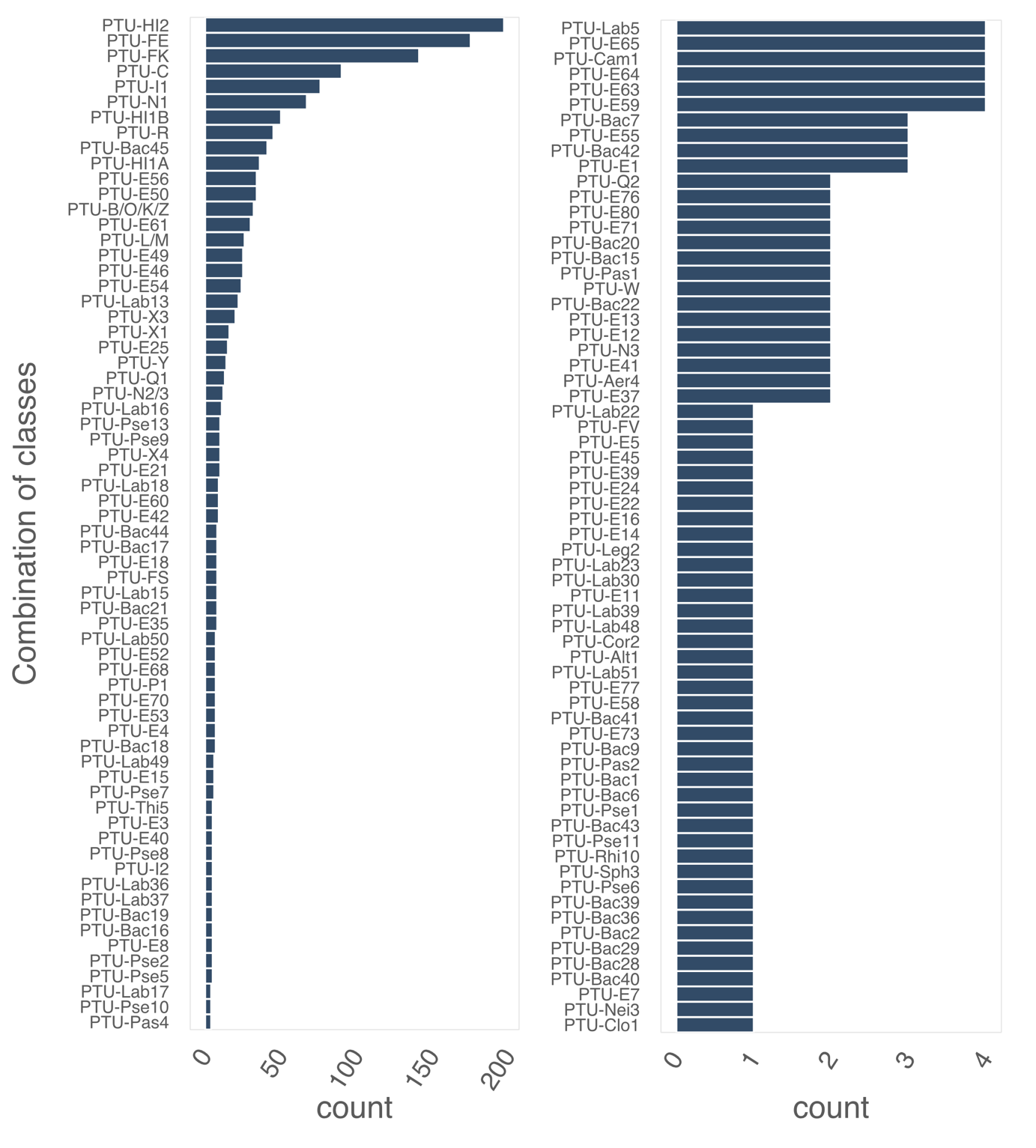

**Figure S3: Number of unique combinations of AMR classes per PTU**. To accommodate differences in scale, the right panel is a continuation from the left panel. The left histogram represents PTUs with 4 to 197 unique combinations of AMR classes, while the right panel represents PTUs with 1 to 4 unique combinations of AMR classes.

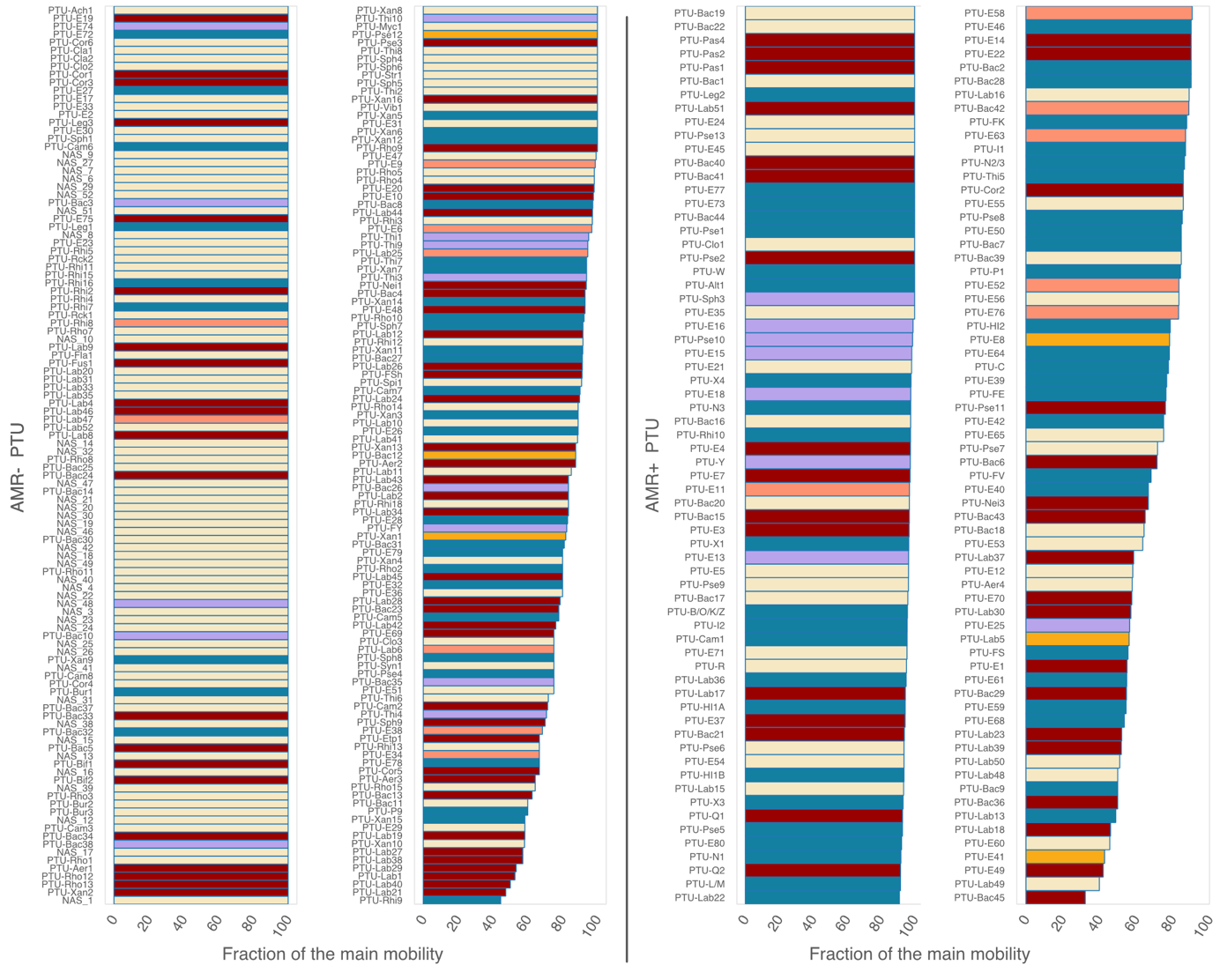
**Figure S4: Proportion of the most abundant mobility type per PTU.** Each PTU was classified according to the most abundant mobility type among the plasmids it contains. The bars represent the frequency of this major mobility type in each PTU. The left panel represents AMR- PTUs, and the right panel represents AMR+ PTUs. In each panel the left histogram is the continuation of the right histogram.

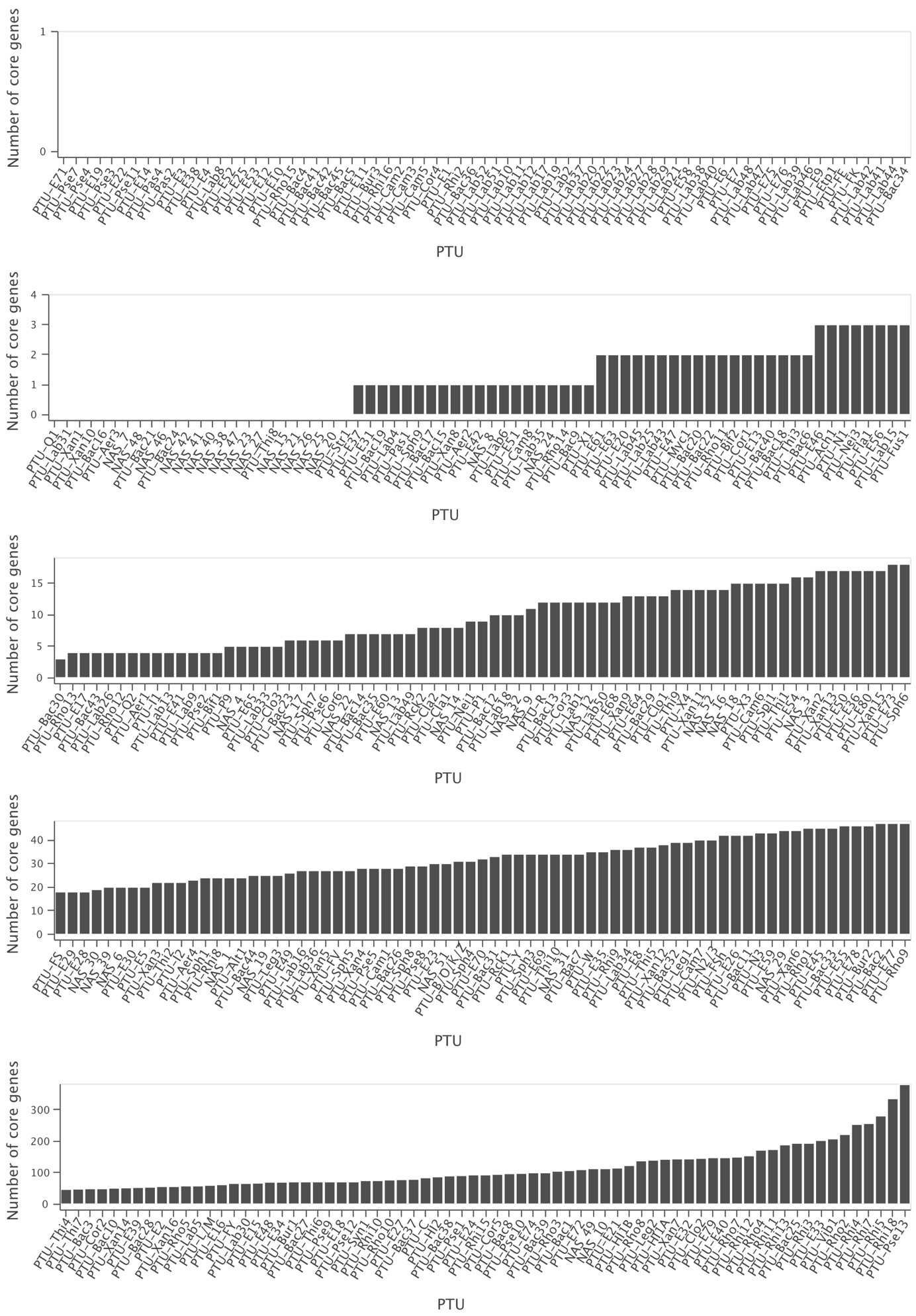
**Figure S5: Distribution of the number of core genes per PTU.** To accommodate differences in scale, each row is the continuation of the previous one.

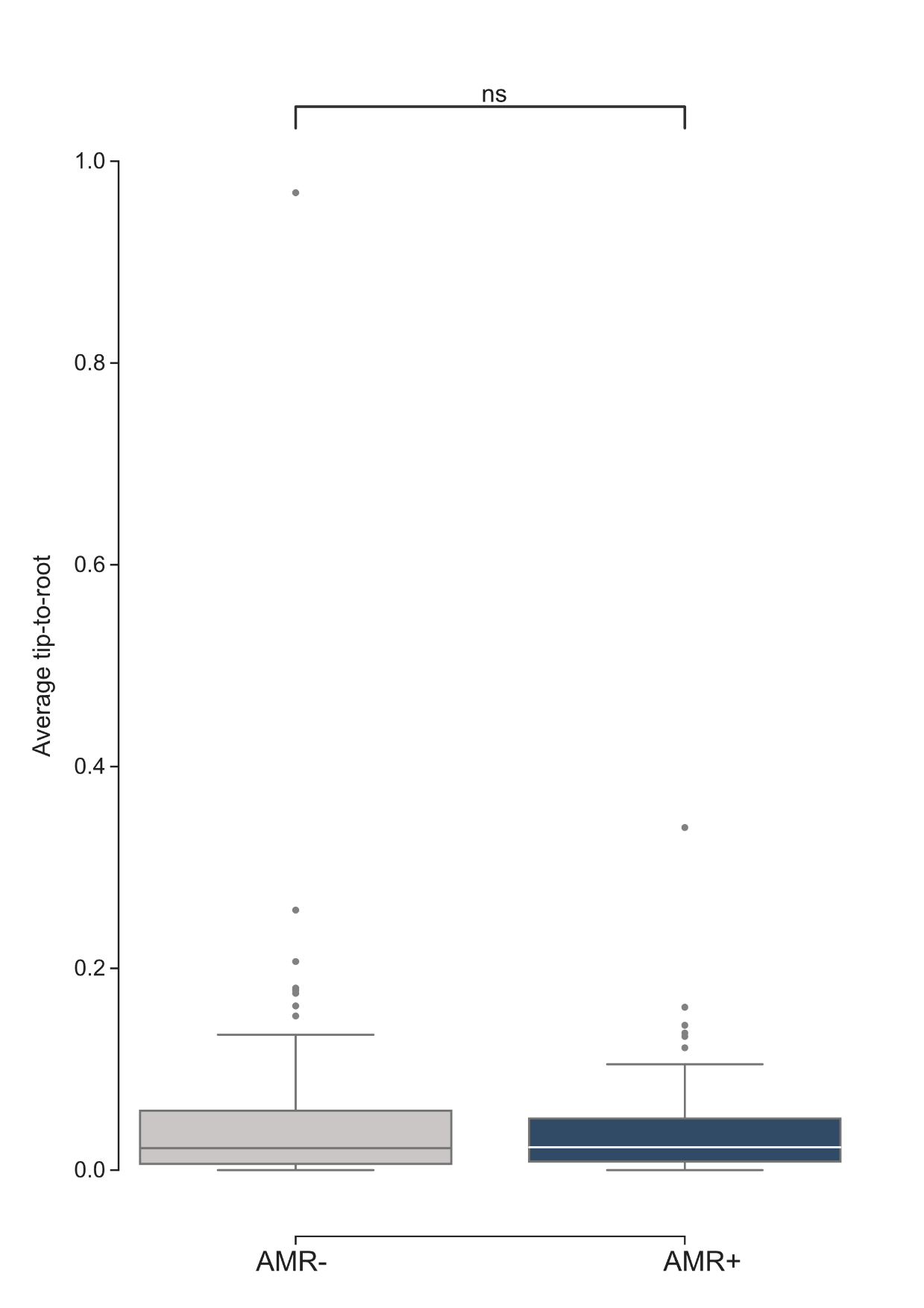

**Figure S6: Distribution of the average tips to root distance between the core genome trees of AMR+ and AMR- PTUs.** The horizontal bar represents the median value, while the lower and upper hinges correspond to the first and third quartiles. The whiskers extend from the hinge to 1.5 times the range between the first and third quartile. Data beyond these values are shown as dots. The differences are not statistically significant (ns: non-significant, P>0.05, Mann-Whitney U).

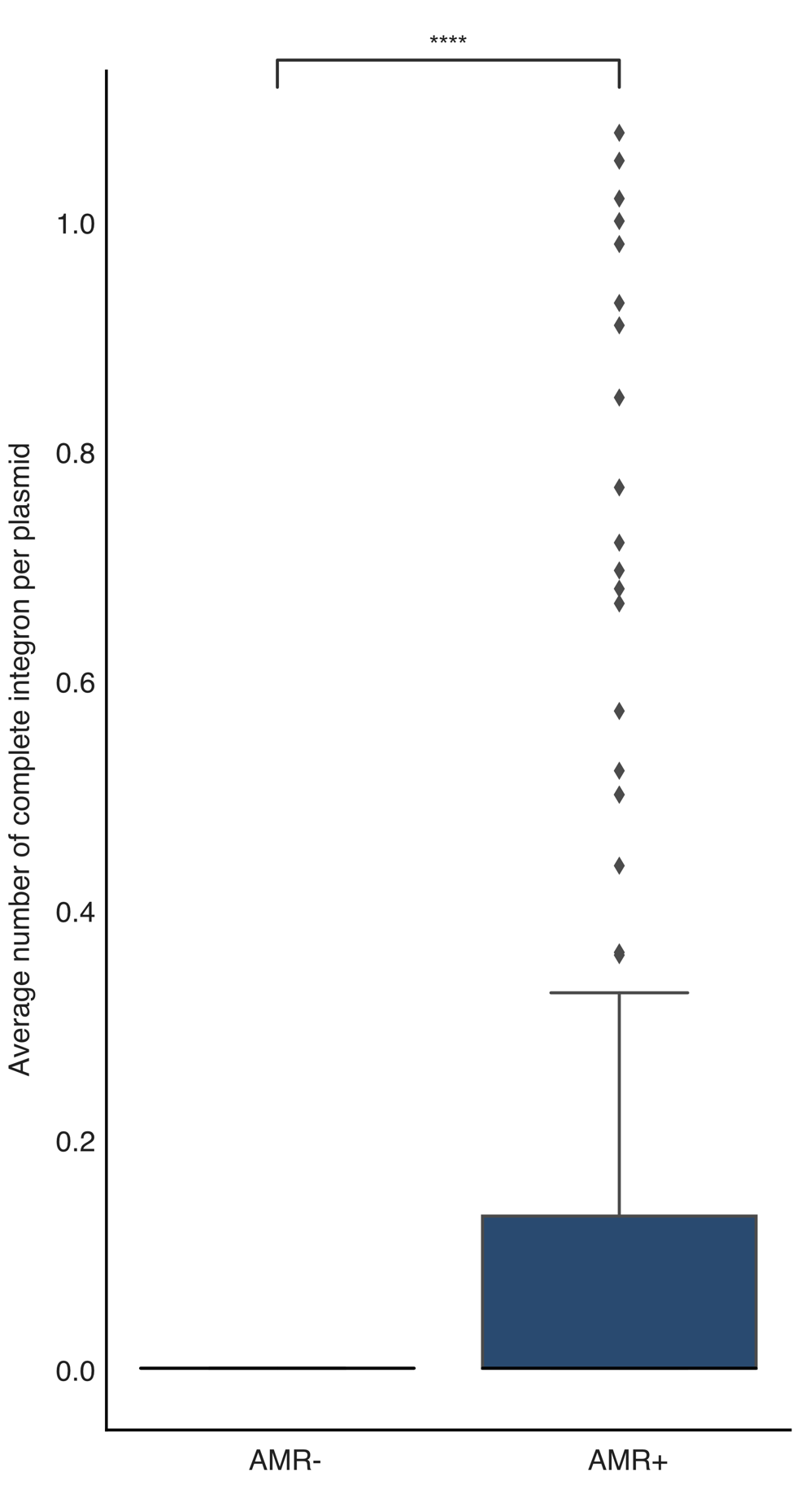

**Figure S7: Distribution of the average number of complete integrons per plasmid between AMR+ and AMR- PTUs.** The horizontal bar in boxplots indicates the median value, and lower and upper hinges correspond to the first and third quartiles. The whiskers extend from the hinge to 1.5 times the range between the first and third quartile. Data beyond these values are shown as dots. Statistical significance (Mann-Whitney U): **** P ≤ 0.0001.
